# Metagenomic Insights Unveil the Dominance of Undescribed Actinobacteria in Pond Ecosystem of an Indian Shrine

**DOI:** 10.1101/475269

**Authors:** Radhakrishnan Manikkam, Madangchanok Imchen, Manigundan Kaari, Angamuthu Vignesh, Venugopal Gopikrishnan, Thangavel Shanmugasundaram, Jerrine Joseph, Ramasamy Balagurunathan, Ranjith Kumavath

**Author notes:** Corresponding author: Dr. M. Radhakrishnan, M.Sc., M.Phil., Ph.D., Scientist E / Associate Professor *Co-Corresponding author: Dr. Ranjith Kumavath Ph.D., FSAB. Equally contributed (Radhakrishnan Manikkam, Ranjith Kumavath).

## Abstract

Metagenomic analysis holds immense potential for identifying rare and uncharacterized microorganisms from many ecological habitats. Actinobacteria have been proved to be an excellent source of novel antibiotics for several decades. The present study was designed to delineate and understand the bacterial diversity with special focus on Actinobacteria from pond sediment collected from Sanjeeviraya Hanuman Temple, Ayyangarkulam, Kanchipuram, Tamil Nadu, India. The sediment had an average temperature (25.32%), pH (7.13), salinity (0.960 mmhos/cm) and high organic content (10.7%) posing minimal stress on growth condition of the microbial community. Subsequent molecular manipulations, sequencing and bioinformatics analysis of V3 and V4 region of 16S rRNA metagenomics analysis confirmed the presence of 40 phyla, 100 classes, 223 orders, 319 families and 308 genera in the sediment sample dominated by Acidobacteria (18.14%), Proteobacteria (15.13%), Chloroflexi (12.34), Actinobacteria (10.84%), Cyanobacteria (5.58%), Verrucomicrobia (3.37%), Firmicutes (2.28%), and, Gemmatimonadetes (1.63%). Among the Actinobacteria phylum, *Acidothermus* (29.68%) was the predominant genus followed by *Actinospica* (17.65%), *Streptomyces* (14.64%), *Nocardia* (4.55%) and *Sinomonas* (2.9%). Culture-dependent isolation of Actinobacteria yielded all strains of similar morphology to that of *Streptomyces* genus which clearly indicating that the traditional based technique is incapable of isolating majority of the non-Streptomyces or the so called rare Actinobacteria. Although Actinobacteria were among the dominant phylum, a close look at the species level indicated that only 15.2% within the Actinobacterial phylum could be assigned to cultured species. This leaves a vast majority of the Actinobacterial species yet to be explored with possible novel metabolites have special pharmaceutical and industrial application. It also indicates that the microbial ecology of pond sediment is neglected fields which need attention.

## INTRODUCTION

It is estimated that only 1% of microbes in the natural environment are culturable under laboratory conditions [46]. More than 88% of culturable isolates belong to four phyla known as *Proteobacteria, Firmicutes, Actinobacteria,* and *Bacteroidetes* [47]. However, uncultivated bacteria possess a great diversity of enzymes adaptable to different environmental conditions which could serve as a prolific source of novel bioactive compounds with industrial and pharmaceutical applications. Metagenomics is a tool to study massive microbial communities prevailing in the environment in its natural state irrespective of its uncultivable properties. This technique has not only helped to overcome the barriers of culture dependent techniques but also identified rare microorganisms [23]. It also provides a wide and unbiased view of the potential microbial functions in a single snapshot with high resolution contributing immensely to the global microbial diversity studies from different ecosystem and their potential ecological functions [27]. On the other hand, there have been several reports on the widespread distribution and occurrence of antibiotic resistance genes [24]. One of the most straightforward ways to tackle the issue is to discover novel antibiotics. *Actinobacteria* are one of the largest phyla ubiquitous to both aquatic and terrestrial ecosystems having high G+C content [48,49] which have produced over 10,000 known antibiotics [15,25]. However, emergences of multidrug resistance have triggered the need for more potent antibiotics. Isolation of rare Actinobacteria could increase the chances of discovering such potentially novel bioactive compounds i.e. Vancomycin was discovered from the rare Actinobacteria strain *Amycolatopsis orientalis* [42, 43]. However, rare Actinobacteria are relatively difficult to isolate and maintain due to the difficulties the lack of appropriate selection media, appropriate isolation procedures and mimicking their natural environment [41]. Antibiotic discovery by conventional approach have the drawback of frequent rediscovery of known metabolites. Hence, focus has to be set on rare microorganisms to have a higher potential of novel metabolites [39, 40, 44]. As Actinobacteria are the well-known antibiotic producers and due to the difficulties of rare Actinobacteria isolation under normal laboratory conditions, metagenomic approach was set to systematically assess the Actinobacterial diversity with respect to the overall microbial structure from pond sediment.

## MATERIALS AND METHODS

### Sampling and physico-chemical characterization of sediment

Sediment samples were collected from the oldest pond of Sanjeeviraya Hanuman Temple, Ayyangarkulam, Kanchipuram district, Tamil Nadu, India (Latitude 79.67′N Longitude 12.78′E) in the month of January, 2018. The sediment samples were pooled and submitted to Omega Laboratories (Analytical Testing and Research Centre, Tamil Nadu, India) for physico-chemical analysis.

### DNA extraction and next generation sequencing

Total metagenomic DNA was extracted from freshly collected sediment sample using PowerWater® DNA Isolation Kits (MoBio Laboratories) following manufacturer′s protocol. DNA quantification was performed using a Qubit™ 3.0 Fluorimeter (Life Technology Ltd., Paisley, UK) and verified by gel electrophoresing on 2% agarose gel. The 260/280 ratio was measured using a Biophotometer (Eppendorf, Hamburg, Germany). DNA samples were stored at −20°C until sequencing was performed. Samples were sequenced at the UAMS Sequencing Core Facility. Bacterial16S rRNA gene V3 and V4 regions were amplified using primers containing Illumina adapters following Illumina’s 16Smetagenomics protocol. Briefly, Kapa Library Amplification Kit was used for PCR and products were cleaned using Beckman Coulter Agencourt AMPure XP Beads according to the16S Metagenomics protocol. The universal 16s forward and reverse primer sequences (**Supplementary data 1**) were implemented to create a single amplicon of approximately 250bp. Concentrations were adjusted to 4uM and prepared for loading on the Illumina Miseq according to Illumina’s 16S metagenomics protocol. Samples were pooled, denatured, and loaded on the Illumina Miseq at 8pM and sequenced paired end (2 × 300) using a MiSeq® Reagent Kitv3.

### Bioinformatics and Statistical Analysis

QIIME 1.9.1 [31] pipeline was used for the entire downstream analysis. Quality check was performed using FastQC0.11.7 [36] and PHRED score reads with >Q30 were considered for further analysis. High quality reads were adapter trimmed and the paired-end reads were stitched using FLASH 1.2.11 [32] to make consensus FASTA sequences. Consensus reads were formed with 0 mismatch having an average contig length of 350 to 450bp and queried to UCHIME [33] to remove all the chimeric sequence which was subsequently pooled and clustered into Operational Taxonomic Units (OTUs) based on their sequence similarity using Uclust (similarity cutoff = 0.97) [35]. Representative sequence was identified for each OTU against SILVA OTUs database using PyNAST [34]. We analyze the microbial diversity within the samples by calculating Shannon, Chao1 and observed species metrics. The chao1 metric estimates the species richness while Shannon metric is the measure to estimate observed OTU abundances, and accounts for both richness and evenness. The observed species metric is the count of unique OTUs identified in the sample.

### Culture dependent isolation of Actinobacteria

Isolation and enumeration of Actinobacteria were performed on selective media Starch Casein Agar (SCA) and Kusters Agar (KUA) supplemented with Nalidixic acid 50µg/ml and Nystatin 20µg/ml. Soil sample was serially diluted up to 10^−5^ and one milliliter of the serially diluted samples was spread into above mentioned media and incubated at room temperature for 7 days. Microscopic characteristics such as the presence of aerial mycelium, substrate mycelium and mycelial fragmentation were observed under the bright field microscope at 40x magnification. Cultural characteristics such as growth, colony consistency, aerial mass colour, reverse side pigment and soluble pigment was recorded by growing the culture on ISP2 (International Streptomyces Project Medium-2) agar medium.

### Antimicrobial activity

Antimicrobial activity of Actinobacteria cultures were tested by adopting agar plug method against *Staphylococcus aureus, Bacillus cereus*, *Escherichia coli, Pseudomonas aeruginosa* and *Klebsiella pneumoniae*. The bacterial strains were grown in nutrient agar medium at 37°C. Actinobacterial cultures were grown on YEME (Yeast Extract-Malt Extract) agar plates for 10 days at 28°C. Bacterial pathogens were spread on LB (Luria bertani) modified agar plate using sterile cotton swab. Agar plug with 5mm diameter which contains the secreted Actinobacteria metabolites were cut from the YEME agar medium in sterile condition and placed over LB modified agar seeded with test pathogens. All the plates were incubated at 37ºC for 24 hours. Zone of inhibition was measured after incubation and expressed in millimetre in diameter.

## RESULTS

### Pond sediment physico-chemical properties

The physico-chemical property of the studied sediment sample is summarized in **Table 1**. The overall physico-chemical properties of the sample was moderate with an electrical conductivity (EC) and total salinity of the sediment of 0.952 mmhos/cm and 0.960 mmhos/cm respectively and the water content of the sediment at the sampling moments was medium. Although calcium and zinc were the dominant cation in sample, the sampling area showed to be dynamic with respect to salinity contributions from different anions and cations from one year to the other. We observed a neutral pH (7.13) and higher organic matter (10.7%) content in the sample retrieved in January 2018.

**Table 1.** Physico-chemical characteristics of sediment sample AKTS

| **Test parameter** | **Unit** | **Result** |
| --- | --- | --- |
| *Color* |  | Red |
| *pH* |  | 7.13 |
| *Salinity* | Mmhos/cm | 0.960 |
| *Acidity* | Mg/kg | 5.82 |
| *Humidity* | g/m<sup>3</sup> | 24.10 |
| *Water activity* |  | Medium |
| *Temperature* | °C | 25.32 |
| *Soil moisture* | % | 24.55 |
| *Silt* | % | 55.6 |
| *Clay* | % | 25.2 |
| *Sand* | % | 18.4 |
| *Electrical conductivity* | Mmhos/cm | 0.952 |
| *Permeability* | Mm/hr | 26.34 |
| *Organic matter* | % | 10.7 |
| *Nitrogen* | % | 0.9 |
| *Phosphate* | % | 2.1 |
| *Potassium* | % | 0.86 |
| *Lime status* | % | 5.8 |
| *Magnesium* | mg/lit | 220 |
| *Sodium* | mg/lit | 145 |
| *Calcium* | mg/lit | 280 |
| *Zinc* | mg/lit | 264 |
| *Copper* | mg/lit | 22 |
| *Boron* | mg/lit | 12 |
| *Aluminum* | mg/lit | 14 |
| *Chromium* | mg/lit | 12 |
| *Fluoride* | mg/lit | 8 |
| *Silver* | mg/lit | 2 |
| *Mercury* | mg/lit | 2 |
| *Selenium* | mg/lit | 4 |

### Microbial community structure

The AKTS-1 microbiota resulted in 514249 high-quality reads with an average read length of 250 bp and 55.09 % total GC content. Base quality score of each cycle in more than 80% of the total reads had phred score greater than 30 (>Q30; error-probability >=0.001). We observed 513710 reads passed mismatch filter which were further processed for removal of chimera and singletons. Rarefaction curve indicated that a reasonable number of individuals were sampled (**Figure 1A-C**). All the resulting fragments were classified into 40 phyla, 100 classes, 223 orders, 319 families and 308 genera. The represented phyla were dominated by unknown phylum (26.8%) followed by Acidobacteria (18.14%), Proteobacteria (15.13%), Chloroflexi (12.34), Actinobacteria (10.84%), Cyanobacteria (5.58%), Verrucomicrobia (3.37%), Firmicutes (2.28%), and, Gemmatimonadetes (1.63%) (**Figure 2A**). Further affiliation revealed that the most abundant genus was affiliated to phylum Actinobacteria which included genus Acidothermus (3.21%), Actinospica (1.91%) and Streptomyces (1.58%). Remaining major genera included *Candidatus solibacter*(3.05%) and *Candidatus koribacter*(1.32%) of Acidobacteria phylum, *Anaerolinea* (2.74%) of Chloroflexi phylum, *Bradyrhizobium* (1.7%) of Proteobacteria phylum. However, most of the genuses were unidentified and uncultured (**Figure 3**). Within the Actinobacteria phyla, although the uncultured (5.28%) and unclassified (17.84%) OTUs were high, it was dominated by several known genus such as Acidothermus (29.68%), Actinospica (17.65%), Streptomyces (14.64%), Uncultured Actinomycete (5.28%), Nocardia (4.55%), Sinomonas (2.90%), Catenulispora (2.57%), Geodermatophilus (1.60%), and, Mycobacterium (0.95%) (**Figure 2B**).

**Figure 1.**
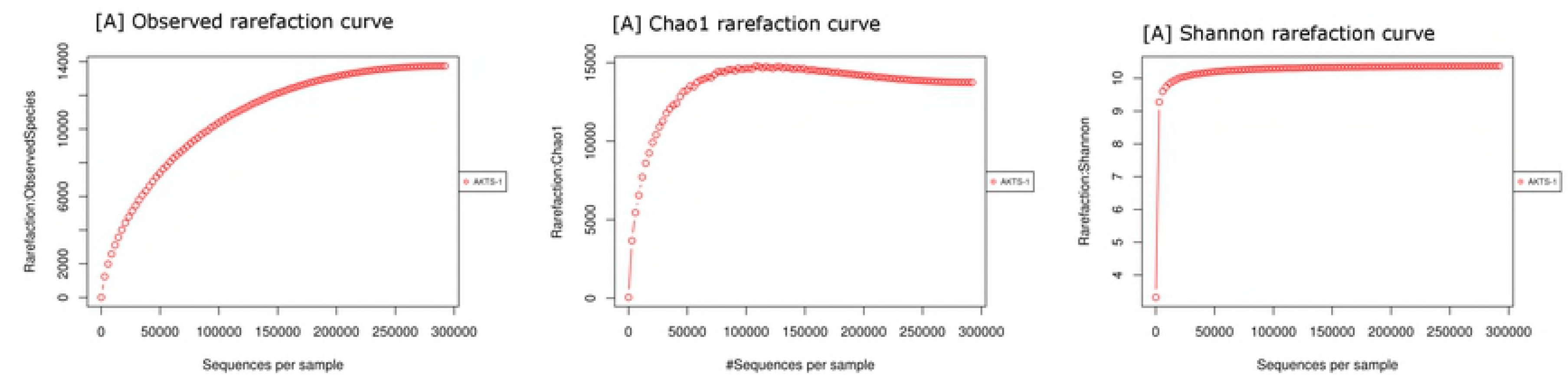
Rarefaction plot of the phylogenetic diversity, using [A] observed [B] chao1 and [C] Shannon curve as alpha diversity metric. The curves reached a plateau indicating the high depth sequencing of the sample.

**Figure 2.**
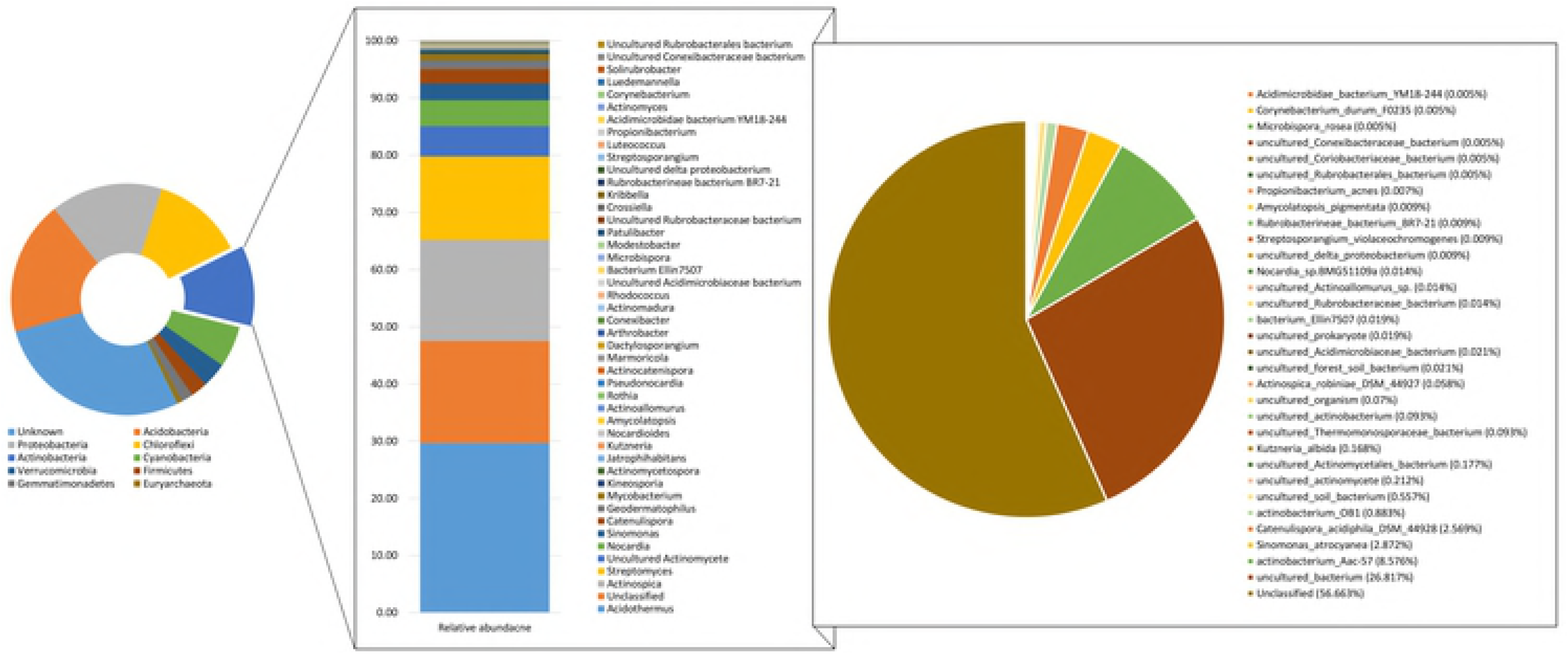
Stacked bar and dough nut chart representing [A] dominant prokaryotic phylum and [B] the list of all Actinobacterial genus and [C] species.

**Figure 3.**
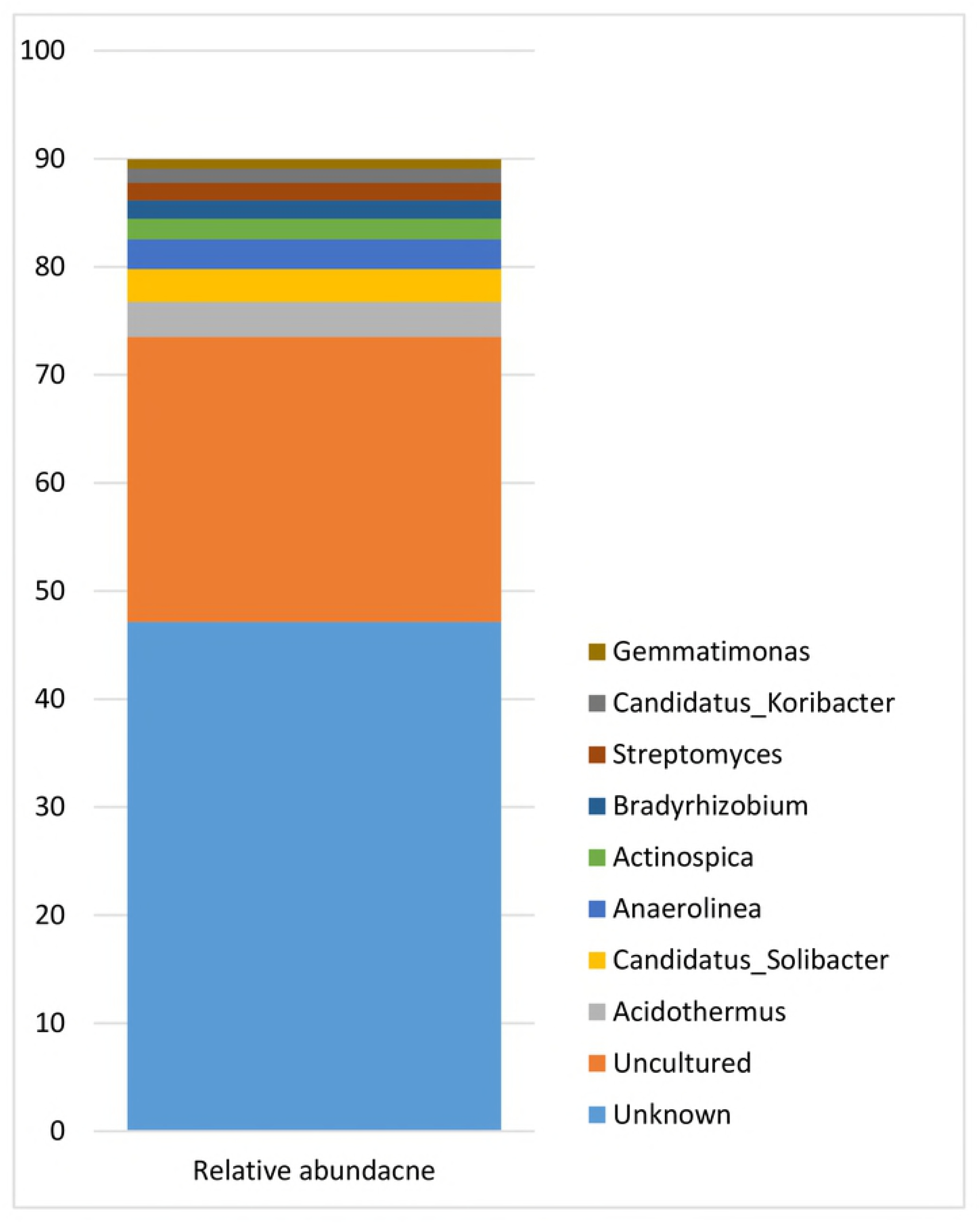
Summary of the top 10 genus predominant in the pond sediment.

### Isolation and antimicrobial screening of Actinobacterial strains

Eighteen morphologically different Actinobacterial strains were isolated from sediment sample (**Table 2**). On ISP2 agar medium, Actinobacterial strains produced powdery growth (n 10) whereas the remaining eight strains produced rough (n 5) and leather (n 3) growth. All the cultures showed the presence of aerial and substrate mycelium under bright field microscopic observation confirming that *Streptomyces* genus dominated in this isolation. Further, in agar plug method, nine out of 18 Actinobacterial strains showed inhibition against at least one of the five clinical pathogens tested. Notably, strains AKTS3 and AKTS11were found to be active against all the five tested pathogens (**Table 2**).

**Table 2.** Details of Actinobacterialstrains isolated from sediment sample and their antimicrobial activity

| Strain | Cultural characteristics |  |  |  |  |  |  | Antimicrobial activity (mm) |  |  |  |  |
| --- | --- | --- | --- | --- | --- | --- | --- | --- | --- | --- | --- | --- |
|  | Growth | Consistency | AMC | RSP | SP | AM | SM | *E. coil* | *Bacillus* | *S. aureus* | *K. pneumonia* | *Candida* |
| AKTS-1 | Good | Powdery | White | Brown | + | + | + |  | 14 |  |  |  |
| AKTS-2 | Good | Powdery | Orange | Pale yellow | - | + | + |  |  |  |  | 9 |
| AKTS-3 | Good | Powdery | Pink | Pale red | - | + | + | 19 | 25 | 15 | 12 | 20 |
| AKTS-4 | Good | Powdery | Grey | Brown | - | + | + |  |  |  |  |  |
| AKTS-5 | Good | Powdery | Dirty white | Red | - | + | + |  |  |  |  |  |
| AKTS-6 | Good | Rough | Orange | Reddish brown | - | + | + |  |  |  |  |  |
| AKTS-7 | Good | Powdery | White | pink | + | + | + |  |  |  |  |  |
| AKTS-8 | Good | Rough | White | - | - | + | + |  |  |  |  |  |
| AKTS-9 | Good | Rough | Pale yellow | Pale yellow | - | + | + |  |  | 28 |  |  |
| AKTS-10 | Good | Powdery | Pale yellow | Pale yellow | - | + | + |  |  |  |  |  |
| AKTS-11 | Good | Leathery | Orange | Pale yellow | - | + | + | 16 | 17 | 30 | 11 | 14 |
| AKTS-12 | Good | Powdery | Pale yellow | Pale red | - | + | + |  |  |  |  |  |
| AKTS-13 | Good | Leathery | Grey | Pale red | - | + | + |  |  |  |  |  |
| AKTS-14 | Good | Leathery | Grey | Pale yellow | - | + | + |  |  |  |  |  |
| AKTS-15 | Good | Rough | Grey | Pale yellow | - | + | + |  | 23 |  |  |  |
| AKTS-16 | Good | Powdery | White | Brown | + | + | + |  | 23 | 18 |  |  |
| AKTS-17 | Good | Powdery | Dirty white | Pale red | - | + | + |  | 16 | 11 |  | 18 |
| AKTS-18 | Good | Rough | Dirty white | Light green | - | + | + |  | 18 |  |  | 11 |

## DISCUSSION

Freshwater bodies account merely 1% of the total water on earth with uneven distribution around the globe. However, freshwater is necessary for majority of plants, animals and for our survival. Microbes play a major role in maintaining water health; its diversity can also indicate a number of parameters regarding water health. Hence, understanding the microbial diversity of the ponds is important. Temple ponds in India serve as a reservoir of freshwater for religious rituals as well as a source for drinking, bathing, washing and daily use. Such ponds are also exposed to various anthropogenic and natural activities giving rise to different carbon sources. Previous studies have found that the water quality as well as microbial community of pond is effected by carbon source [19]. In natural ecosystems, physiochemical conditions such as pH, soils type, temperature, salinity and other organic contents have high impact on metabolic activity and distribution of microbiota [20]. The presence of average electrical conductivity, salinity, and, ion concentration in the sediment sample within the study indicated positive microbial activity and a minimal stress on the microbial community.

### Pond sediment microbial community structure

Community structure of pond sediment is known to be extremely complex and only a few dominant phyla are capable of thriving under such severely competitive environmental conditions. Hence, the dominant bacterial phyla in pond sediment were identified and explored as a part of this study. Dominant phyla included Acidobacteria (18.14%), Proteobacteria (15.13%), Chloroflexi (12.34%) and Actinobacteria (10.84%) (**Figure 2A**). However, a major part of the reads were assigned to unknown phylum (26.8%). Previous reports have reported contrasting dominance of phylum in pond sediments. Actinobacteria were seen to be the most dominated phylum in freshwater agricultural pond, mine area pit pond, rainforest soil and Freshwater lake surface [1, 6, 14, 17]. In case of aquaculture pond sediments and ponds located at the vicinity to hexachlorocyclohexane (HCH) production, Proteobacteria were the most dominant [16, 19]. However, in this report, Acidobacteria was the most dominant phylum in freshwater pond sediment. Although the race for the most dominate phylum might not show a clear pattern, it is interesting to note that Actinobacteria stands among one of the most abundant phylum in the entire freshwater pond. Interestingly, the most abundant genus *Acidothermus* (3.21%) fell under phylum Actinobacteria which is rather unusual since Acidothermus was originally isolated from Yellowstone National Park acidic hot spring [7]. Acidothermus is a monospecic genus described as thermophilic, acidophilic, and cellulolytic bacterium which can also degrade chitin [2, 3]. However, there are previous reports of genus Acidothermus being enriched in the salinized rhizosphere of *Tamarixparviflora* [4]. It was also observed that the pond sediment exhibited 308 genera (**Supplementary data 2**) out of which 46.9% of the genus were unclassified indicating the potential novel genus yet to be discovered. In addition, 26.39% of the genus were uncultured. This shows that more than 70% of the reads mapped to genera have never been insolated in pure culture. The isolation of microbes in its pure culture form is of outmost importance in order to study its ecological roles and the potential array of metabolites that can have huge industrial and pharmaceutical applications.

### Actinobacterial community

Actinobacterial genus such as *Acidothermus* (3.21%), *Actinospica* (1.91%), and, *Streptomyces* (1.58%) dominated the bacterial genera indicating that the Actinobacterial genus are a dominated member of bacterial Kingdom in pond sediment (**Figure 3**). These genera are also found abundantly in various soil specimens and are possibly enriched in the pond sediment due to presence of aromatic hydrocarbons and other similar factors [16]. A closer look at genus level specifically within the Actinobacterial community highlighted eight predominant genera namely *Acidothermus* (29.6%), *Actinospica* (17.65%), *Streptomyces* (14.64%), *Nocardia* (4.55%), *Sinomonas* (2.90%), *Catenulispora* (2.57%) and *Geodermatophilus* (1.6%) dominated in varying proportions throughout the study period (**Figure 2B**). This is in sharp contrast to freshwater river, lake and agricultural pond being dominated by *Actinomycetales* and *Streptomyces* [17, 10]. Although Streptomyces was not the most dominated in this study, it occupied the 3^rd^ most abundant genus within Actinobacterial phylum indicating its predominant nature. Interestingly, Streptomyces is the largest genus within Actinobacterial phylum and accounts for over 75% of the bioactive compounds production from Actinobacterial phylum [8]. Within species level, a vast majority of the reads (abundance) were either unclassified (56.6%) or uncultured (28.13%) (**Figure 2C**). This is a clear indication that a large section of Actinobacterial species are unexplored and untapped of its full potential. The large proportion of Actinobacterial phylum within bacterial kingdom, however, with substantial unclassified OTUs at species level also suggests that those undescribed bacterial groups with unknown metabolic capabilities are an important part of the pond microbiota, a matter which warrants further investigation. In fact, looking at the Actinobacterial species diversity, only 15 out of 32 OTUs were classified into known species (**Figure 2C**) accounting for merely 15.2% of the Actinobacterial abundance which indicates the dominance of yet-to-be cultured Actinobacterial species. Since majority of microbes are not culturable under laboratory conditions, it is no wonder that there have been an exponential decline of novel bioactive compounds discovery from Actinobacteria [9].

Actinobacteria as a promising reservoir of potential novel antibiotics is far from exhausted [46] and research has been extensively performed from academic to industrial groups in quest of novel metabolites [29]. Actinobacteria is a major reservoir of bioactive compounds housing over 45% of all microbial origin. Hence, in this study we have isolated 18 Actinobacterial strains in pure culture using SCA selective media. Half of the isolated colonies showed antimicrobial activity against at least one of the five different clinical pathogens tested. Strains AKTS3 and AKTS11 exhibited antimicrobial activity against all the five different clinical pathogens tested (**Table 2**). It is apparent that the discovery of novel rare Actinobacteria can be expected to provide new bioactive compounds [26, 37, 42].

In the past years, focus on natural product discovery from Actinobacteria shifted from extensively investigated soil-dwelling strains towards underexplored habitats of rare Actinobacteria from unusual ecosystems [30]. This approach can give a significant impact on the discovery platform for novel compounds with promising bioactivities from rare Actinobacteria [45]. However, it is clear that the soil dwelling Actinobacteria from freshwater pond harbors a significant amount of unexplored Actinobacterial species. There were also several bacterial rare species, below 0.001%, (n 186; **Supplementary data 3**) of a great diversity with low abundance, 16 belonging to either unknown or uncultured Actinobacteria phylum. It has been suggested that members of the rare biosphere may play important roles in the bacterial community [22, 28]. Freshwater ecosystem, especially stagnant water bodies such as lakes have shown some promising results for rare Actinobacterial genus [10, 17]. Development of new selective techniques and expression based metagenomics would enable screening, isolation, and discoveries of rare antibiotic genes which could lead to useful bioactive compounds.

## CONCLUSION

The study shows the presence of unexplored rare and uncultured Actinobacteria through metagenomics. The work also clearly demonstrates that most of the actinobacteria strains have high antimicrobial activity. However, the current traditional selective media is not suitable for harvesting rare and uncultured Actinobacteria community. Hence, potential novel secondary metabolite from rare Actinobacteria, especially from exotic environmental habitats, is still underexplored. Focus has to be highlighted to develop novel insolation strategies and media to isolate such rare microbes. Neglecting the rare isolates would understate the immense potential of novel bioactive compounds and their role as a keystone species in the environment.

## Author contributions

VG; MK performed the experiments, RM conceived and designed the experiments, analyzed the data, contributed reagents/materials/analysis tools, prepared figures and/or tables, Wrote the paper: RK RM, MI, MK, AV, VG, TS, JJ, RB, All authors gave critical advice on the manuscript and approved the version to be published.

## Acknowledgments

Authors acknowledge the Sathyabama Institute of Science, Technology and Central University of Kerala for the research facilities provided. The authors also acknowledge the financial support from the Department of Science & Technology-Science and Engineering Research Board, SERB- (No.SB/051/2014) Govt. of India.

## Ethical approval

This article does not contain any studies with pathogenic organisms or anthropogenic activities performed by any of the authors.

## Data availability

All data relevant to the article is included in the article and its supplementary information.

## REFERENCES

1. Lima-Perim JE, Romagnoli EM, Dini-Andreote F, Durrer A, Dias ACF, & Andreote FD (2016). Linking the composition of bacterial and archaeal communities to characteristics of soil and flora composition in the Atlantic Rainforest. PloS one, 11(1), e0146566.

2. Lynch MD, & Neufeld JD (2016). SSUnique: Detecting Sequence Novelty in Microbiome Surveys. MSystems, 1(6), e00133-16.

3. Barabote RD, Xie G, Leu DH, Normand P, Necsulea A, Daubin V et al. (2009). Complete genome of the cellulolytic thermophile Acidothermus cellulolyticus 11B provides insights into its ecophysiological and evolutionary adaptations. Genome Res 19: 1033–1043.

4. Polivkova M, Suman J, Strejcek M, Kracmarova M, Hradilova M, Filipova A, & Uhlik O (2018). Diversity of root-associated microbial populations of Tamarix parviflora cultivated under various conditions. Applied Soil Ecology, 125, 264-272.

5. Berry AM, Barabote RD, & Normand P (2014). The Family Acidothermaceae. In The Prokaryotes (pp. 13-19). Springer, Berlin, Heidelberg.

6. Monard C, Gantner S, Bertilsson S, Hallin S, & Stenlid J (2016). Habitat generalists and specialists in microbial communities across a terrestrial-freshwater gradient. Scientific reports, 6, 37719.

7. Mohagheghi A, Grohmann KMMH, Himmel M, Leighton L, & Updegraff DM (1986). Isolation and characterization of Acidothermus cellulolyticus gen. nov., sp. nov., a new genus of thermophilic, acidophilic, cellulolytic bacteria. International Journal of Systematic and Evolutionary Microbiology, 36(3), 435-443.

8. Berdy J. (2005) Bioactive microbial metabolites. J Antibiot. 58:1–26.

9. Alvan G, Edlund C, Heddini A. (2011) the global need for effective antibiotics-summary of plenary presentations. Drug Resis Updat. 14:70–6.

10. Zothanpuia Passari AK, Chandra P, Leo VV, Mishra VK, Kumar B, et al. (2017) Production of potent antimicrobial compounds from Streptomyces cyaneofuscatus associated with fresh water sediment. Front Microbiol. 8:68.

11. Ningthoujam DS, Sanasam S, Nimaichand S. (2011) Studies on bioactive actinomycetes in a Niche Biotope, Nambul River in Manipur, India. J Microbial Biochem Technol. S6:001.

12. Sanasam S, Nimaichand S, Ningthoujam D. (2011) Novel bioactive actinomycetes from a niche biotope, Loktak Lake, in Manipur, India. J Pharm Res. 4:1707–10.

13. Jami M, Ghanbaria M, Kneifela W, Domig KJ. (2015) Phylogenetic diversity and biological activity of culturable actinobacteria isolated from freshwater fish gut microbiota. Microbiol Res. 175:6–15

14. Driscoll HE, Vincent JJ, English EL, & Dolci ED. (2016) Metagenomic investigation of the microbial diversity in a chrysotile asbestos mine pit pond, Lowell, Vermont, USA. Genomics data, 10, 158-164.

15. Radhakrishnan M, Jerrine J, Anbarasu S, Suresh A, and Balagurunathan R. (2017). Actinobacteria from less explored ecosystems: A promising source for anti TB metabolites. (Book Chapter). In: Current Research in Microbiology (Online), USA. 1-29. (ISBN: 978-93-87500-01-3)

16. Negi V, & Lal R. (2017) Metagenomic Analysis of a Complex Community Present in Pond Sediment. Journal of genomics, 5, 36-47

17. Chopyk J, Allard S, Nasko D, Bui A, Mongodin EF, & Sapkota AR. (2018) Agricultural freshwater pond supports diverse and dynamic bacterial and viral populations. Frontiers in microbiology, 9, 792.

18. Newton RJ, Jones SE, Eiler A, Mcmahon KD, Bertilsson S. (2011) A guide to the natural history of freshwater lake bacteria. Microbiol. Mol. Biol. Rev. 75 14–49. 10.1128/M

19. Qin Y, Hou J, Deng M., Liu, Q, Wu, C, Ji, Y, & He X. (2016) Bacterial abundance and diversity in pond water supplied with different feeds. Scientific reports, 6, 35232.

20. Anjum R, Malik A. Evaluation of mutagenicity of wastewater in the vicinity of pesticide industry. Environ ToxicolPharmacol. 2013; 35: 284–291.

21. Ara I, Bakir MA, Hozzein WN, and Kudo T. (2013). Population, morphological and chemotaxonomical characterization of diverse rare actinomycetes in the mangrove and medicinal plant rhizosphere. Afr.J. Microbiol.Res. 7, 1480–1488. doi:10.5897/AJMR12.452

22. Bagatini IL, Eiler A, Bertilsson S, Klaveness D, Tessarolli LP, Vieira AA. Host-specificity and dynamics in bacterial communities associated with Bloom-forming freshwater phytoplankton. PloS One. 2014; 9(1):e85950. doi:10.1371/journal.pone.0085950 PMID: 24465807

23. Lin KH, Liao BY, Chang HW, Huang SW, Chang TY, Yang CY, … & Yu HT. (2015) Metabolic characteristics of dominant microbes and key rare species from an acidic hot spring in Taiwan revealed by metagenomics. BMC genomics, 16(1), 1029.

24. Imchen M, Kumavath R, Barh D, Vaz A, Góes-Neto A, Tiwari S, …& Azevedo V. (2018) Comparative mangrove metagenome reveals global prevalence of heavy metals and antibiotic resistome across different ecosystems. Scientific reports, 8(1), 11187.

25. Moopantakath J, & Kumavath R. (2018). Bio-Augmentation of Actinobacteria and Their Role in Dye Decolorization. In New and Future Developments in Microbial Biotechnology and Bioengineering (pp. 297-304).

26. Bérdy J. (2005) Bioactive microbial metabolites (review). J. Antibiot. 58, 1–26. doi: 10.1038/ja.2005.1

27. Brady SF, Simmons L, Kim JH, Schmidt EW. (2009) Metagenomic approaches to natural products from free-living and symbiotic organisms. Nat. Prod. Rep. 26, 1488–1503.

28. Cao Y, Williams DD, Williams NE. How important are rare species in aquatic community ecology and bioassessment? LimnolOceanogr. 1998; 43(7):1403-9.

29. Genilloud O. (2017) Actinomycetes still a source of novel antibiotics. Nat. Prod. Rep. 34, 1203–1232.

30. Balagurunathan R, Radhakrishnan M, and Ramya S. (2014) Extremophilic and extremotolerant Actinomycetes: Distribution and Importance. In: Recent Trends in Microbial Diversity and Bioprospecting. Westville Publishing House, New Delhi (ISBN: 978-93-83491-14-8); pp. 114-128.

31. Kuczynski J, Stombaugh J, Walters WA, González A, Caporaso JG, & Knight R. (2012). Using QIIME to analyze 16S rRNA gene sequences from microbial communities. Current protocols in microbiology, 27(1), 1E-5.

32. Magoč T, & Salzberg SL. (2011) FLASH: fast length adjustment of short reads to improve genome assemblies. Bioinformatics, 27(21), 2957-2963.

33. Edgar RC, Haas BJ, Clemente JC, Quince C, & Knight R. (2011) UCHIME improves sensitivity and speed of chimera detection. Bioinformatics, 27(16), 2194-2200.

34. Caporaso JG, Bittinger K, Bushman FD, DeSantis TZ, Andersen GL, & Knight R. (2009) PyNAST: a flexible tool for aligning sequences to a template alignment. Bioinformatics, 26(2), 266-267.’

35. Edgar RC. (2010). Search and clustering orders of magnitude faster than BLAST. Bioinformatics, 26(19), 2460-2461.

36. Andrews S. (2010). FastQC: a quality control tool for high throughput sequence data. Available online at: http://www.bioinformatics.babraham.ac.uk/projects/fastqc

37. Hong K, Gao AH, Xie QY, Gao H, Zhuang L, Lin HP, et al. (2009) Actinobacteria for marine drug discovery isolated from mangrove soils and plants in China. Mar. Drugs 7, 24–44. doi:10.3390/md70 10024

38. Hubert J, Nuzillard JM, Renault, JH. (2017) Dereplication strategies in natural product research: How many tools and methodologies behind the same concept? Phytochem. Rev. 16, 55–95.

39. Hug JJ, Bader CD, Remškar M, Cirnski K, Müller R. (2018) Concepts and Methods to Access Novel Antibiotics from Actinomycetes. Antibiotics 7, 44; doi:10.3390/antibiotics7020044

40. Katz L, Baltz RH. (2016) Natural product discovery: Past, present, and future. J. Ind. Microbiol. Biotechnol. 43, 155–176.

41. Khanna M, Solanki R, and Lal R. (2011). Selective isolation of rare actinomycetes producing novel antimicrobial compounds. Int. J. Adv. Biotechnol. Res. 2, 357–375.

42. Lee LH, Zainal N, Azman AS, Eng SK, Goh BH, Yin et al. (2014). Diversity and antimicrobial activities of actinobacteria isolated from tropical mangrove sediments in Malaysia. Scientific World J. 2014:698178. doi:10.1155/2014/698178.

43. Mahajan GB, and Balachandran L. (2012) Antibacterial agents from actinomycetes-are view. Front. Biosci. 4:e373. doi:10.2741/E373

44. Penesyan A, Kjelleberg S, Egan S. (2010) Development of novel drugs from marine surface associated microorganisms. Mar. Drugs 8, 438–459.

45. Tiwari K, and Gupta RK. (2012) Rare actinomycetes: a potential store house for novel antibiotics. Crit. Rev. Biotechnol. 32,108–132. doi:10.3109/07388551.2011.562482

46. Vartoukian SR, Palmer RM, Wade WG. (2010) Strategies for culture of “Cunculturable” bacteria. FEMS Microbiol. Lett. 309, 1–7.

47. Nikolaki S, Tsiamis G. (2013) Microbial diversity in the era of omic technologies. BioMed. Res. Int. 2013:958719. 10.1155/2013/958719.

48. Ventura M, Canchaya C, Tauch A, Chandra G, Fitzgerald GF, Chater KF, & van Sinderen D. (2007) Genomics of Actinobacteria: tracing the evolutionary history of an ancient phylum. Microbiology and molecular biology reviews: MMBR, 71(3), 495-548.

49. Barka EA, Vatsa P, Sanchez L, Gaveau-Vaillant N, Jacquard C, et al. (2016) Taxonomy, physiology, and natural products of Actinobacteria. Microbiol Mol Biol Rev. 80(1):1–43.

